# Molecular Learning of a Soft-Disks Fluid

**DOI:** 10.1101/2021.07.24.453642

**Authors:** Luca Zammataro

**Affiliations:** Kiromic Biopharma

**Keywords:** molecular learning, molecular computing, molecular dynamics, thermodynamics, machine learning, Lennard-Jones fluid, self-organizing systems, logic-gates

## Abstract

This work is based on the Equivalence between Molecular Dynamics and Neural Network. It provides learning proofs in a Lennard-Jones (LJ) fluid, presented as a network of particles having non-bonded interactions. I describe the fluid’s learning as the property of an order that emerges as an adaptation in establishing equilibrium with energy and thermal conservation. The experimental section demonstrates the fluid can be trained with logic-gates patterns. The work goes beyond Molecular Computing’s application, explaining how this model uses its intrinsic minimizing properties in learning and predicting outputs. Finally, it gives hints for a theory on real chemistry’s computational universality.

## 1 Introduction

Molecular Computing is a promising scientific frontier with its origin as early as 1961, thanks to Feynman (Adleman 1994). It started to be concretely developed in the mid-1980s, with Conrad (Sienko 2003), and successively in the mid-1990s with Adelman (Adleman 1994). Studies of these authors have demonstrated that biological substrates can be used for finding solutions to combinatorial problems at a molecular level (Adleman 1994). Molecular Computing aims to construct information processing systems in which individual molecules play a functional role. Some examples encompass materials able to emulate biology or follow *de novo* architectural principles, imitating conventional silicon architectures like wires, switches, and “nano-islands.” These are molecular systems that, for the Coulomb effect, can encode bits in the presence or absence of one electron (Sienko 2003).

Maybe, Molecular Computing researchers focused more on computing based on DNA nanotechnology than small synthetic molecules or peptides able to process information. The reason is that DNA computing can quickly treat problems requiring parallelizable tasks. Thus, DNA strands can hold a large amount of data in memory and drive multiple operations at once.

Anyway, even though the baggage of knowledge around DNA computing nowadays reached a considerable level, it is not clear whether it is possible to create circuitries able to learn without making a complex system involving many DNA strands.

Moreover, supposing we imagine a possible involvement of single molecules in molecular learning, we may have to face difficulty understanding how a single molecule can implement an information processing system through its internal structure. We may blame this difficulty for its entropy.

The age-old experience of Molecular Dynamics (MD) can help identify the learning mechanisms in molecule systems. MD algorithms developed in the last forty years aims to answer how the order can locally emerge at the expense of an increase in external entropy (Nicolis and Prigogine 1977). It is no coincidence that these algorithms share strong similarities with Machine Learning.

The formal analogy between thermodynamics and learning has a long story (Gori, Maggini et al. 2016, Goldt and Seifert 2017, Hylton 2020). The literature describes several models that link thermodynamics to the conservation of information (the first law) and the relative entropy decrease (second law). These models range over from disordered lattice Ising-spin-glasses to Markovian processes and Neural Networks (Hylton 2020). In these last, kinetic and potential energy concepts interpret learning as a dissipation-driven adaptation mechanism. In other words, it is a self-emergent regularization process that works on a training set’s approximation accuracy (Gori, Maggini et al. 2016, Hylton 2020).

Therefore, in Neural Networks, dynamics leading learning can be described as a dissipative process, in which the kinetic energy reflects the weight variations. In contrast, the potential energy represents the penalty relating to the degree of satisfaction of the environmental constraints. (Gori, Maggini et al. 2016, Hylton 2020).

In Machine Learning, the goal is to minimize the distance between the real and the predicted output, represented by the standard error and embodied in the logistic regression cost function J(θ), where θ correspond to weights. The cost function is given by:

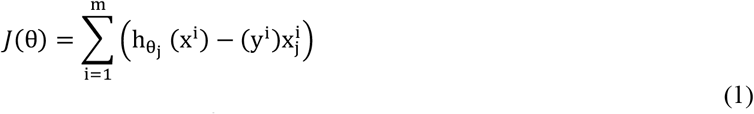

Where *h* _*θ*_, the hypothesis, is the matrix product of *θ*^*T*^*x*; x^i^is the features vector, *m* is the number of patterns in the training set and y^i^is the actual outcome.

Neural Networks must calculate a cost function that involves *j l*evels of neurons. For a cost function of such magnitude, the searching task for the best θ is considered an optimization problem. For this reason, they are provided by sophisticated functions, like *gradient descent*, which simultaneously updates θ to find *min*_*θ*_*J*(*θ*), by calculating the partial derivative. Therefore, Neural Networks show conservation, dissipation, adaptation, and self-organization (Hylton 2020). Remarkably, all these properties belong to any other network as an open thermodynamic system (Hylton 2020).

This work demonstrates that chemical systems can also show information processing abilities through their particles’ self-organization, intended as building blocks. For example, a fluid can be represented as a network of interconnected elements, whereas a specific potential energy’s function handles their interactions. Therefore, a single fluid particle represents an artificial neuron, while a force field, intended as expressions of potential energies, governs the particle interactions.

The work is based on the Equivalence between Molecular Dynamics and Neural Network, a concept discussed for the first time. It draws inspiration from a two-dimensional soft-disks simulation of a fluid (Rapaport 2004). The cost function here is represented by the Lennard-Jones (LJ) potential, which governs the particle’s pair interactions. The simultaneous minimization of all the potential energies makes the simulation a gradient descent algorithm; that’s why integrating all the forces between atom pairs works like a gradient.

Finally, experiments shed light on the fluid’s learning, and they are described in the Results. For the experiments, I have decided to use patterns of six logic gates (AND, NAND, OR, NOR, XOR, and XNOR) for demonstrating that the soft-disks fluid can be trained with Boolean functions. Logic gates, which are electronic device implementations, can accept inputs and produce outputs based on their state. Moreover, they can be connected to make different switching functions and combinational circuits. Computers are based on them to transform the 1s and 0s from input wires. Logic gates represent idealized computation models, and for this reason, they are a good test for demonstrating computational universality.

The fluid’s learning emerges by the sequential administration of all the logic gates patterns, exploiting the fluid minimization property. This phenomenon could be not limited to the only LJ fluid, and it could involve other chemical systems, like rigid molecules governed by different kinds of potentials.

The work suggests hints for a theory on real chemistry’s computational universality.

## 2 Molecular Learning Model

In the following paragraphs, I will describe a machine with learning abilities based on LJ fluid-dynamics. The learning is possible thanks to a couple of particles’ solicitation defined as the system’s input and one particle as output. For its similarity with Neural Networks, I will introduce the assumption of Equivalence between Molecular Dynamics and Neural Networks. A description of the model implementation will follow.

### 2.1 Lennard-Jones potential

A Lennard-Jones (LJ) fluid is a soft-disks simulation in which the potential, 4ε[(σ/r)^12^ − (σ/r)^6^], models the strength of the interaction and repulsion for a pair of atoms *i* and *j* located at *r*_*i*_ and *r*_*j*_ positions in an *n*-dimensional box (Rapaport 2004). The potential serves for calculating forces between pair of particles separated by *r*, where *r* represents the separation distance between the two atoms in Å units. The parameter σ is a length scale, meaning the distance at which the intermolecular potential between the two atoms is = 0 (in Å units). The formula comprises two components: the repulsive (σ/r^12^) and the attractive term (σ/r^6^), respectively, denoting Pauli repulsion due to overlapping electron orbitals at short ranges, and attractive forces at long ranges (van der Waals force) between pairs of particles. The ε parameter defines the strength of the interaction (in eV units); in essence, ε is a measure of how strongly two particles attract each other. Finally, *u* represents the intermolecular potential between the two particles. The interaction repels at close range, then attracts, and is cut off at some limiting separation *r*_*c*_. As *r* increases towards *r*_*c*_, the force drops to 0. The limiting separation is given by 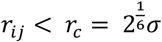. Figure 1 reports a graphical representation of the LJ potential.

**Figure 1.**
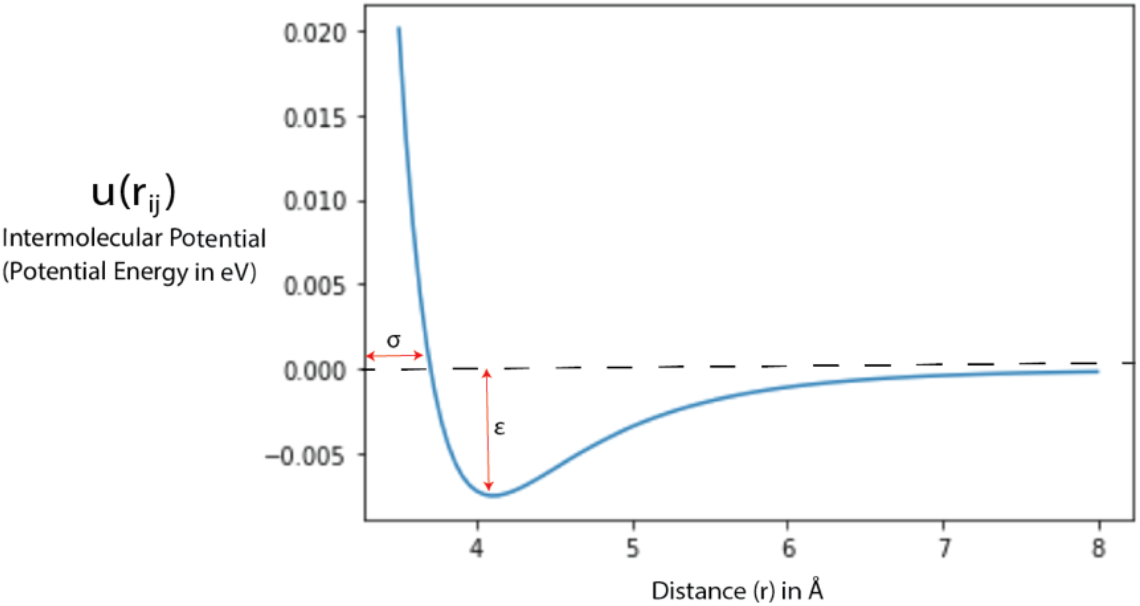
The Lennard-Jones potential. The potential (blue line) is calculated for a couple of particles separated by *r*, representing the separation distance between the two atoms in Å units. The parameter σ is a length scale, that is the distance at which the intermolecular potential between the two atoms is = 0 (in Å units).

### 2.2 Assumption of Equivalence between Molecular Dynamics and Neural Networks

A network of particles has to be considered as a Neural Network. An atom corresponds to a node; its velocity represents the node’s value, and coordinates and accelerations correspond to weights/synapses. The machine’s input (predictor variables) corresponds to a set of atoms embedded in the same system, which undergo a continuous and controlled solicitation of velocity values. The output neuron (response variable) is represented by another particle situated in the same network. During the training phase, the output undergoes a steady solicitation of velocity values. Instead, during the testing, it will be released to provide prediction values. Thus, the particle network is programmable by accessing the input/output (I/O) particle’s weights and modifying the vectors’ velocity values. The essence of Equivalence is summarized in Table 1.

**Table 1.**
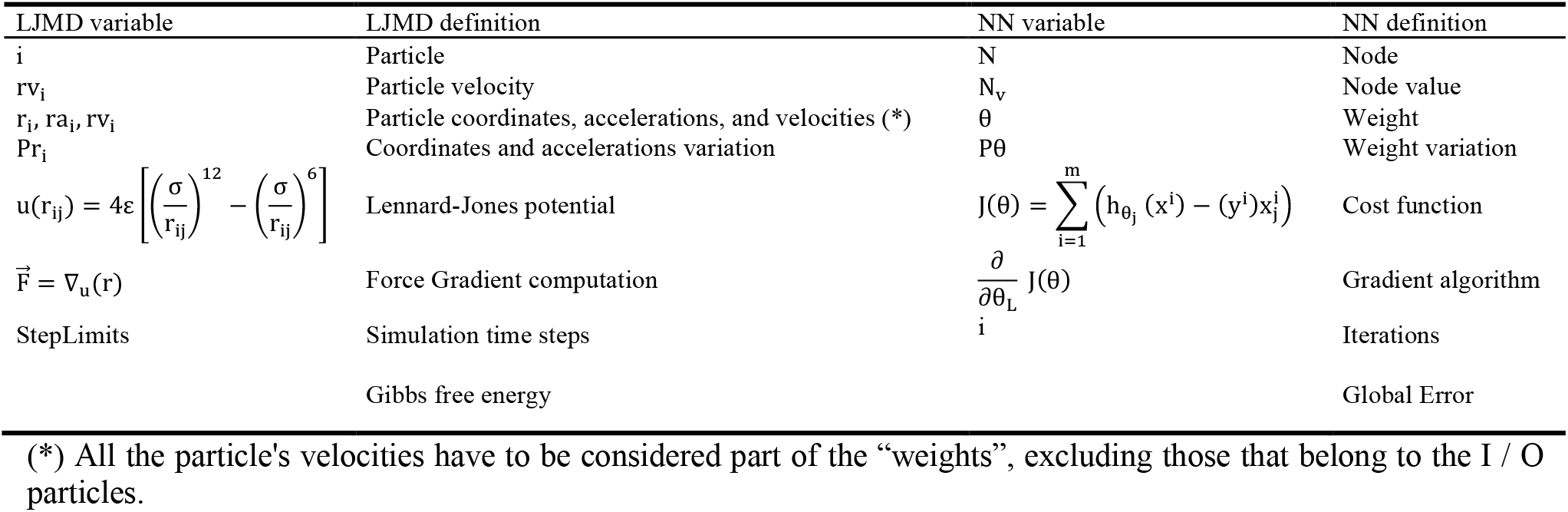
Assumption of Equivalence between Lennard-Jones Molecular Dynamics (LJMD) and Neural Networks (NN).

As a consequence of the Equivalence, a fluid can be described as a machine that generates output for a specific input, evolving in a feature space ℝ^feat^, as expressed by a function of time T: [t_0_, t_1_] → ℝ^feat^. This function of time is then mapped to the output z ∈ ℝ^out^.

Like in a Neural Network, we distinguish a training phase from a testing phase. During the training, the simulation learns I/O patterns by maintaining solicitations of value velocities for the particles designed to cover input and output roles. Instead, during the testing phase, the fluid predicts an output given an input (see paragraph 2.5). Each of these phases is characterized by time steps whose limit must be defined before the program running.

The Equivalence assumes that the LJ potential is comparable to a Neural Network’s cost function, while the force corresponds to the gradient computation. The model works in a two-dimensional box; thus, a two-dimensional vector represents the force that, for each atom pair, is equal and opposite to the potential energy gradient:

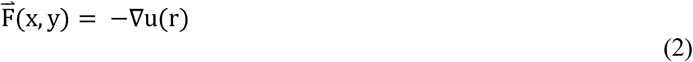

Rewriting (2), we get that a conservative force is equal to the derivate of its potential energy:

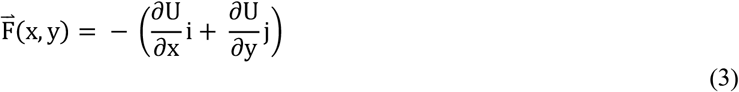

Therefore, each time step, the force calculated for an *ij* atoms pair tends to minimize that pair’s potential. At the end of the training phase, the algorithm produces lists of weight vectors representing the system’s configuration and, at the same time, its knowledge base. The testing phase will use these weights to make predictions given a specific input.

### 2.3. Model Implementation

The Soft-Disks Fluid Learning algorithm (SDFL) implements a two-dimensional lattice of uniform finite elements representing atoms, whose states evolve in a discrete-time, depending on two functions. The first one, called *ComputeForces*, implements the LJP for calculating the forces. The second is a function based on the *Leapfrog* method, a numerical technique for integrating the motion equation. The latter has good energy conservation properties in integrating the atom’s coordinates and velocities (Rapaport 2004).

Thermodynamically, a particle’s state comprises three vectors, *r, v*, and *a*, representing coordinates, velocity, and acceleration. Each vector consists of two directional values *(x, y)* since the model works in two dimensions. Thus, each variable is expressed as the sum of squares of the vector values. All the atoms of the lattice update their states synchronously. *ComputeForces* and *Leapfrog* calculate the next state of every atom basing on the states of its closest neighbors. The approach used to compute interaction is the “all-pairs” method, which means that all the atoms’ pairs must be analyzed since it is not known in advance which particles interact (Rapaport 2004) (Figure 2A).

**Figure 2.**
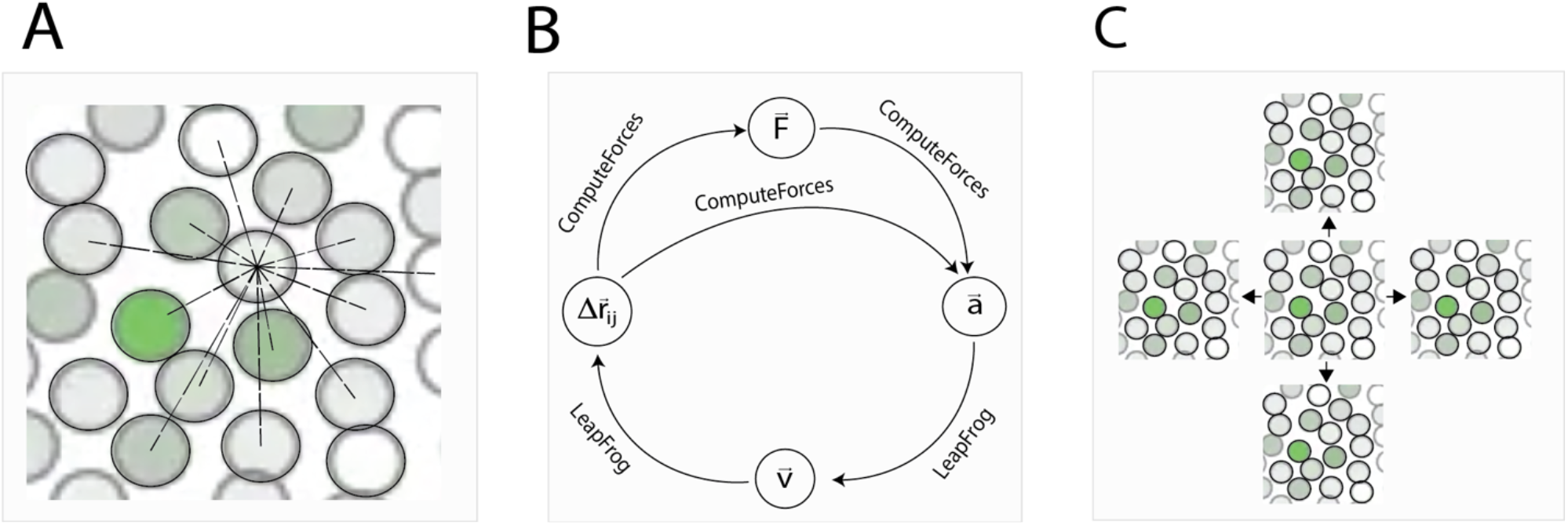
**A**: the approach chosen to compute particle interactions is the “all-pairs” method, which means that all the atoms’ pairs must be analyzed. **B**: schematization of the *ComputeForce* and *LeapFrog* implementation. **C:** periodic boundaries are taken into account to avoid signals losing in the box edges’ proximity.

All the fluid particles’ velocities are set to a fixed magnitude (*velMag*), which depends on the temperature:

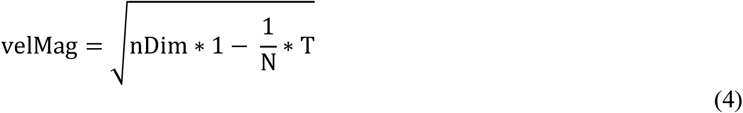

Where *nDim* is the number of dimensions, N the number of particles, and T is the temperature. After randomly assigning velocity directions, the velocities are adjusted to ensure that the mass center is stationary. Each atom pair needs only be examined once. *ComputeForces* calculates the intermolecular potential u(r_ij_) between the two particles through the LJP and synchronously computes the forces applying Newton’s second law, which implies that the force F for each pair of particles at an *r* position is:

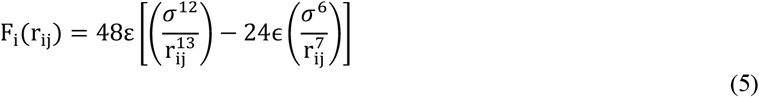

Also, *ComputeForces* updates the accelerations multiplying F_ji_for the difference of a pair of particle coordinates Δr_ji_= r_i_− r_j_. For Newton’s third law, F_ji_= −F_ji_; thus, the acceleration for a couple of particles *ij* will be (Rapaport 2004):

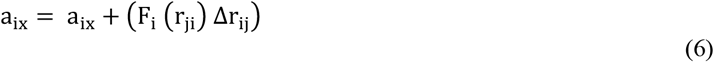

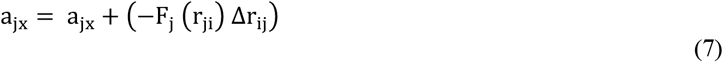

Before *ComputeForces* computes the accelerations, *Leapfrog* updates all the velocities by a half time step, using the old acceleration values (Rapaport 2002):

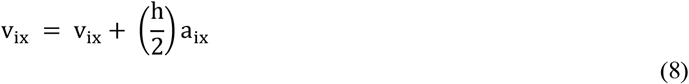

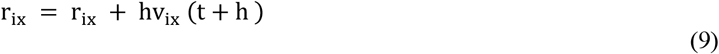

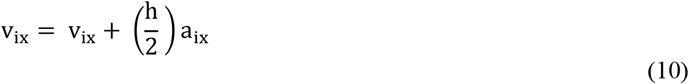

The coordinates are then updated by a full timestep using the intermediate velocity values (Rapaport 2004). A schematization of the model implementation is provided in Figure 2B. Periodic boundaries are taken into account to avoid signals losing in the box edges’ proximity (Figure 2C). The dimensionless kinetic and potential energies for each particle are given by:

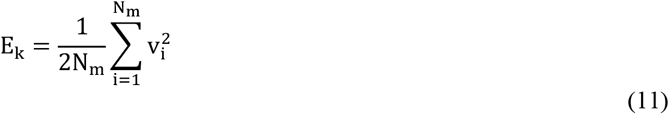

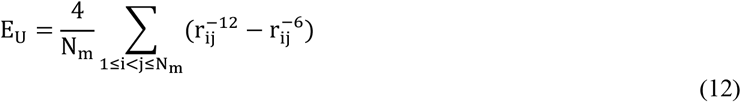

The temperature is expressed in ϵ/Kb

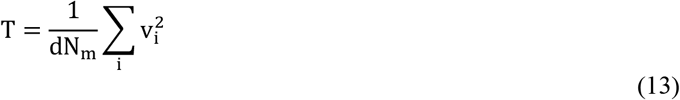

For each time step, global kinetic energy, temperature, and pressure are calculated.

The SDFL algorithm draws inspiration from a two-dimensional soft-disks simulation of a fluid presented by Dennis C. Rapaport in his book “The Art of Molecular Dynamics Simulation” (Rapaport 2004). The code is a Python implementation of the primary functions initially written in C language (Zammataro 2020).

I have introduced one Python class for the molecule definition (class Mol) and one for the properties (class Prop), represented by two C structs, in the original Rapaport’s version. I have replaced all the vector operations functions with linear algebra functions from *NumPy* (Virtanen, Gommers et al. 2020). Also, I have rewritten all the necessary functions for randomness and updating the coordinates in periodic boundaries.

The program comprises the Main loop, the *SetupJob* function, and the *SingleStep* function. Each one represents a wrap of additional subfunctions.

The *SetParams* function serves to set global parameters. At the same time, *SetupJob* embodies *InitCoords, InitVels*, and *InitAccels*, which are functions for the initialization of the coordinates, the velocities, and the accelerations of all the atoms, respectively.

The Main loop works in two modalities: training and testing. In both cases, it calls *SingleStep*, which represents the function that handles the whole process. *SingleStep*, in turn, will call *ComputeForces* and *LeapFrog* aforementioned described. *LeapFrog* appears twice in the listing of *SingleStep*, with the argument 1 or 2 that determines which portion of the two-step leapfrog process has to be performed. *SingleStep* also encompasses two functions for the simulation’s properties measurements: *EvalProps* and *AccumProps*.

In order to demonstrate its learning potentiality, I have used patterns of six logic gates (AND, NAND, OR, NOR, XOR, and XNOR) for training the algorithm with Boolean functions. When the Main loop enters the *training* mode, the *LoadLogicPattern* function uploads a training set, corresponding to a specific logic gate, which comprises four patterns. The training set is processed for some iterations whose number is defined by the parameter *iterations* (Table 4). When the loop ends, it produces lists of weight vectors, saving them as files; in essence, the three files contain coordinates, accelerations, and velocities vectors: they represent the fluid’s knowledge base.

During the *testing* phase, the Main loop uploads the weights and the query files and runs the test making predictions given a specific input pattern. Terminated the *testing*, the output particle O’s velocity values are recorded for *n* timesteps and saved in particular tracks for the analysis. The pseudocode in Figure 3 schematizes the flux diagram of the whole algorithm.

**Figure 3.**
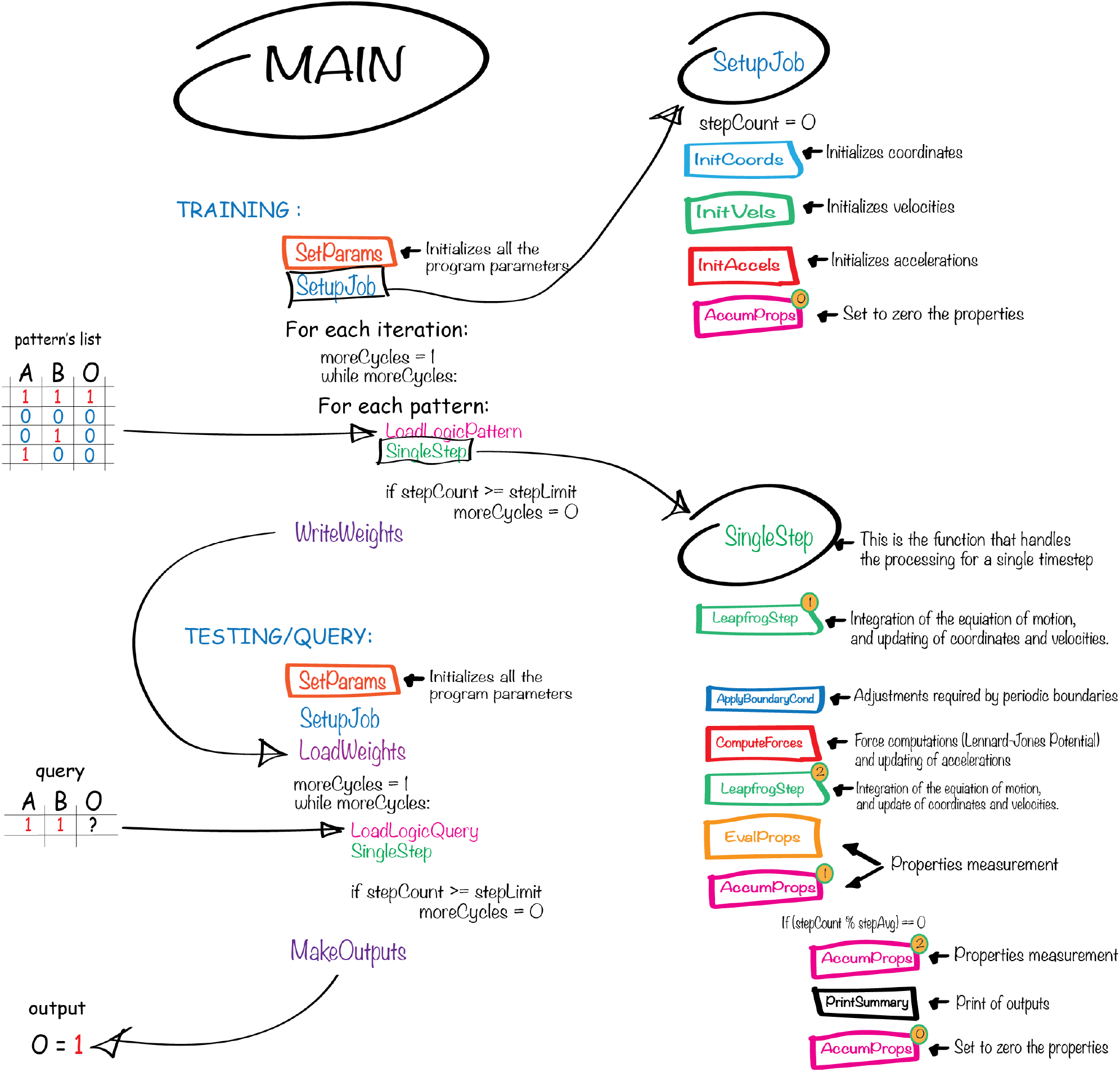
Flux diagram of the Soft-Disks Fluid Learning algorithm (SDFL). The program comprises three parts: The Main loop, the *SetupJob* function, and the *SingleStep* function, which calls *ComputeForces* and *LeapFrog. LeapFrog* appears twice in the listing of *SingleStep*, with the argument 1 or 2 that determines which portion of the two-step leapfrog process has to be performed. *SingleStep* also encompasses two functions for the simulation’s properties measurements: *EvalProps* and *AccumProps*. I.e., *AccumProps* can calculate energy properties or their standard deviations, based on the argument 1 or 2. The Main loop can work in *training* and *testing* mode.

### 2.4. Training

The algorithm has been designed to make the fluid a “programmable system” by training logic-gates patterns. All the parameters are loadable by a couple of parameter files. Each pattern is composed of two input values and one output. During the training phase, the fluid simulation evolves in time. Each pattern is administrated by assigning three on *n* particles; two of them are given to the input (A, B). In contrast, one particle is used for the output O.

As aforementioned said, a particle’s velocity is a vector containing a couple of values *x, y*, (the same for the acceleration vectors). Each value represents velocity directions related to *x* and *y* axes. Thus, the variable is expressed as the sum of squares of the vector values: *rv_x*^*2*^ *+ rv_y*^*2*^. If, for example, we want that the velocity value for an input particle is 9.0, we have to assign the tuple (3.0, 3.0) to its velocity vector. The default velocity value used for the I/O is 2.0 = (1.0, 1.0), for mimicking the Boolean value “true/1”. Contrarily, a velocity equal to 0.0 = (0.0, 0.0) corresponds to the Boolean values “false/0”. I.e., given two input particles A and B and one output particle O, we want to perform training with the AND logic-gate pattern set. We have to solicit the fluid with the AND set reported in Table 2 for all the training phase duration.

**Table 2.** Example of AND logic-gate pattern. The velocity values used for training are tuples representing velocity directions. The variable is expressed as the sum of squares of the tuple.

| Particle Input A | Particle Input B | Particle Output O |
| --- | --- | --- |
| (0.0, 0.0) = 0.0 | (0.0, 0.0) = 0.0 | (0.0, 0.0) = 0.0 |
| (1.0, 1.0) = 2.0 | (0.0, 0.0) = 0.0 | (0.0, 0.0) = 0.0 |
| (0.0, 0.0) = 0.0 | (1.0, 1.0) = 2.0 | (0.0, 0.0) = 0.0 |
| (1.0, 1.0) = 2.0 | (1.0, 1.0) = 2.0 | (1.0, 1.0) = 2.0 |

In contrast, the three (A, B, O) particles’ spatial coordinates are kept to fixed values, preventing their update and avoiding their fluctuation in the two-dimensional box. For accessing the training phase, the algorithm has to upload an eighteen lines file containing simulation settings. The *LoadLogicPattern* function (Figure 3) lists patterns specific for the logic-gate set chosen for the training. An example of the file is described in Table 3.

**Table 3.** Example of a AND logic-gate training file.

|  |  |  |
| --- | --- | --- |
| 1 | deltaT | 0.005 |
| 2 | density | 0.8 |
| 3 | initUcell_x | 20 |
| 4 | initUcell_y | 20 |
| 5 | stepAvg | 100 |
| 6 | stepEquil | 0 |
| 7 | stepLimit | 1000 |
| 8 | temperature | 1 |
| 9 | mode | training |
| 10 | iterations | 10 |
| 11 | movie | no |
| 12 | A_pos | 170 |
| 13 | B_pos | 171 |
| 14 | O_pos | 271 |
| 15 | pattern | 0 0 0 |
| 16 | pattern | 1 1 1 |
| 17 | pattern | 0 1 0 |
| 18 | pattern | 1 0 0 |

Lines from 1 to 8 contain settings for global simulation’s parameters like *initUcell_x and initUcell_y*, which define a 20X20 two-dimensional particle box (400 particles) of mass *μ*. Parameters *σ*, and *ε*, are set to 1 as default, thus they are not listed in the file. Other parameters are *density, temperature*, and *stepLimit*. The density of the fluid is set to 0.8, while the initial temperature is 1. The parameter *stepLimit* sets the number of cycles for the fluid simulation; each cycle is a *while* loop running until the number of cycles exceeds *stepLimit*. The *deltaT* value is related to the energy conservation properties of the *Leapfrog* method implemented here, and aforementioned, discussed in paragraph 2.3. The parameter *stepAvg* controls the number of time steps for printing the output of the molecular dynamics. Lines 12-14 define the I/O particle positions. I.e., A_pos = 170, means that the A input is located at position 170 in the fluid box.

Lines from 9 to 11 contain the necessary information for the simulation. The parameter *training* must be set to “yes” if we want the algorithm performs training and not testing. The parameter *Iterations* plans how many learning iterations the algorithm has to do.

Finally, lines from 15 to 18 that start with the keyword *pattern* define a specific logic-gate pattern for training the fluid. Three values characterize each pattern. I.e., the pattern in line 15 corresponds to the logic: input A=0, input B=0, output=0. One important caveat is that the training is strongly limited to the sequential presentation of the patterns.

### 2.5. Testing/Query

During the testing, the algorithm uses the weights produced in training for making predictions given a specific query. The particles involved in the testing process try to reproduce the output values to fit the training experience. We have to provide a particular file (Table 4) that differs from the training for the *mode* parameter, which has to be set to “testing”. Moreover, instead of *patterns*, we have to provide a *query*. The algorithm’s function preposed for the query upload is *LoadLogicQuery* (Figure 3).

**Table 4.** Example of an AND logic-gate query/testing file: The case of input A=1, B=1.

|  |  |  |
| --- | --- | --- |
| 1 | deltaT | 0.005 |
| 2 | density | 0.8 |
| 3 | initUcell_x | 20 |
| 4 | initUcell_y | 20 |
| 5 | stepAvg | 100 |
| 6 | stepEquil | 0 |
| 7 | stepLimit | 1000 |
| 8 | temperature | 1 |
| 9 | mode | testing |
| 11 | movie | no |
| 12 | A_pos | 170 |
| 13 | B_pos | 171 |
| 14 | O_pos | 271 |
| 15 | query | 1 1 |

I.e., the *query* in line 15 is formed by a couple of values for the two input particles. When we run the testing, we expect that the fluid produces the corresponding output of one of the AND conditions (A=1, B=1), thus O=1.

## 3 Results

The parameters used for all the experiments are: 400 particles distributed in a 20X20 two-dimensional particle box; *σ*, and *ε*, are set to 1; fluid’s density = 0.8, and initial temperature = 1. The I/O particles are assigned to positions 170 for input A, 171 for input B, and 271 for the output O.

The velocity values used for training and testing are set to 2.0 = (1.0, 1.0), for mimicking the Boolean value “true/1”, and 0.0 = (0.0, 0.0) for the Boolean values “false/0”. In one case on six, (the XOR case), the fluid shows better performances in predicting outputs, with velocities values of 1.0 = (0.5, 0.5).

The number of iterations for the training is ten in all the cases, except for the OR gate, which needs only five iterations. The number of timesteps used for training is 1000. When the simulation starts (timesteps=0), all the particles are equidistantly disposed, and velocity and acceleration directions are randomly assigned. The sum of potential energies (Ep) is ∼ 0.66 (Figure 4A).

**Figure 4.**
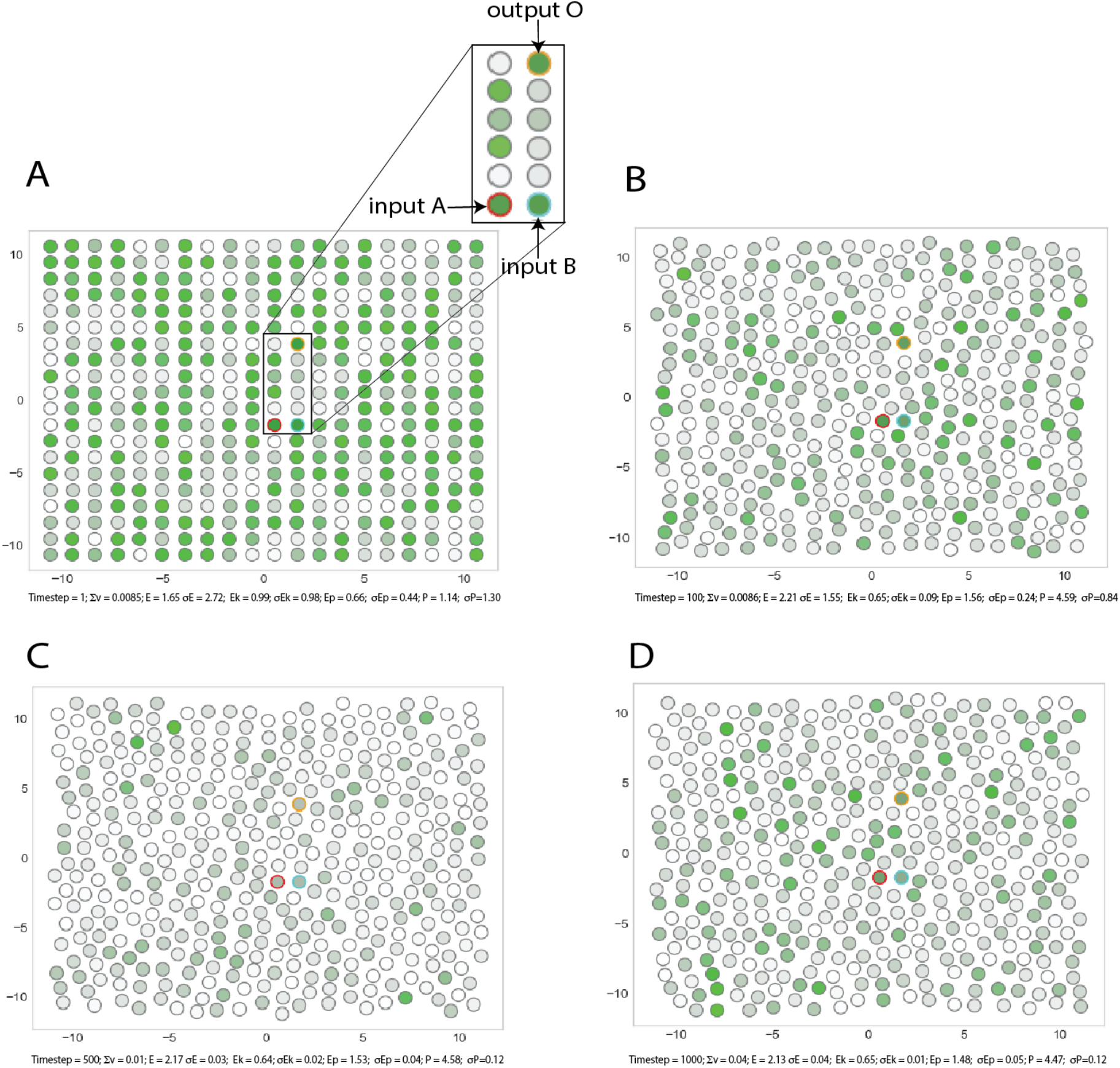
Graphical output of four moments of the SDFL algorithm during the AND training phase. The 400 particles are distributed in a 20X20 two-dimensional box. In the plot, each particle assumes shades of green based on the particle velocity. The green intensity is normalized on the max velocity values among all the particles. The I/O particles are centrally disposed at the positions 170 for A, 171 for B, and 271 for O. Their borders are colored in red, cyan, and yellow, respectively for A, B, and O. **A:** the program starts with all the particles equidistantly positioned. The sum of potential energies (Ep) is ∼ 0.66. **B**: after 100 timesteps, the particles are sparse with non-ordered trajectories. (Ep=1.55). **C:** after 500 timesteps, the minimizing process produces ordered particles’ trajectories in a solid and dense fluid phase (Ep=1.53). **D:** at timestep 1000, the Ep is 1.48.

Take a glance at the AND case: after 100 timesteps, the particles appear to be sparse with non-ordered trajectories, as shown in Figure 4B, and the Ep increases to 1.55. After 500 timesteps, the minimizing process produces ordered particles’ trajectories in a solid and dense fluid phase (Figure 4C). The Ep is reduced to 1.53. When the fluid makes this type of trajectory, the gradient starts its minimizing process; the particles’ distances reach a compromise since the computation of the force selects coordinates that imply minimum costs for fluid, thus minimum potential energy. At timestep 1000, the minimizing process reaches a good performance; the Ep is 1.48 (Figure 4D).

Sum of the global velocities components, mean energy, kinetic energy, potential energy per atom, pressure and standard deviations during the AND training, are reported in Table 5.

**Table 5.**
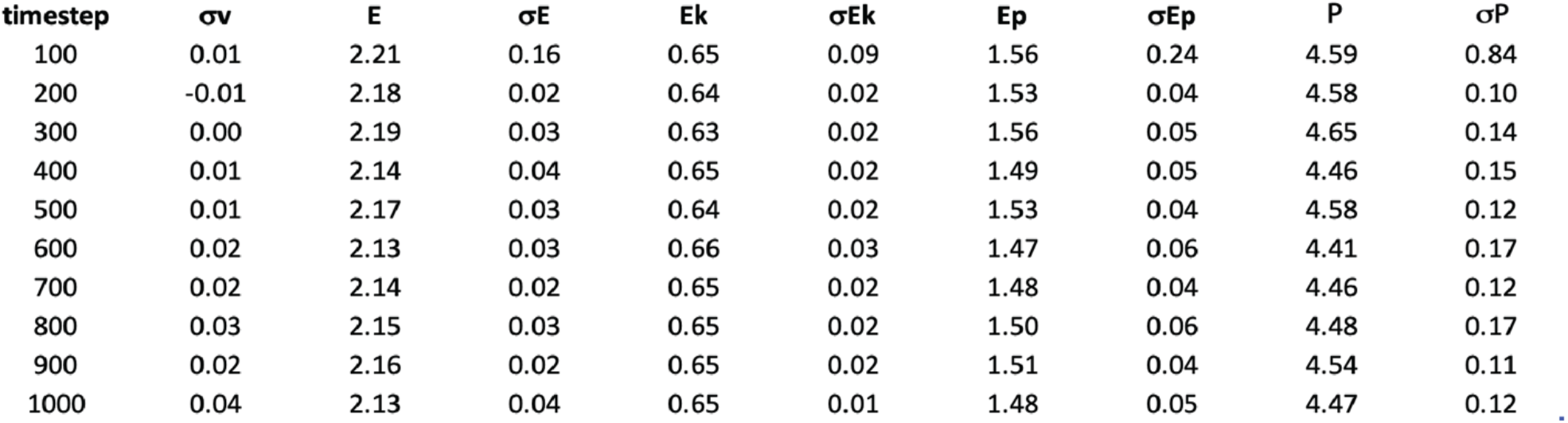
Sum of the global velocities’ components, mean energy, kinetic energy, potential energy per atom, pressure and standard deviations during the AND training.

| timestep | $\sigma v$ | E | $\sigma E$ | $E_k$ | $\sigma E_k$ | $E_p$ | $\sigma E_p$ | P | $\sigma P$ |
| --- | --- | --- | --- | --- | --- | --- | --- | --- | --- |
| 100 | 0.01 | 2.21 | 0.16 | 0.65 | 0.09 | 1.56 | 0.24 | 4.59 | 0.84 |
| 200 | -0.01 | 2.18 | 0.02 | 0.64 | 0.02 | 1.53 | 0.04 | 4.58 | 0.10 |
| 300 | 0.00 | 2.19 | 0.03 | 0.63 | 0.02 | 1.56 | 0.05 | 4.65 | 0.14 |
| 400 | 0.01 | 2.14 | 0.04 | 0.65 | 0.02 | 1.49 | 0.05 | 4.46 | 0.15 |
| 500 | 0.01 | 2.17 | 0.03 | 0.64 | 0.02 | 1.53 | 0.04 | 4.58 | 0.12 |
| 600 | 0.02 | 2.13 | 0.03 | 0.66 | 0.03 | 1.47 | 0.06 | 4.41 | 0.17 |
| 700 | 0.02 | 2.14 | 0.02 | 0.65 | 0.02 | 1.48 | 0.04 | 4.46 | 0.12 |
| 800 | 0.03 | 2.15 | 0.03 | 0.65 | 0.02 | 1.50 | 0.06 | 4.48 | 0.17 |
| 900 | 0.02 | 2.16 | 0.02 | 0.65 | 0.02 | 1.51 | 0.04 | 4.54 | 0.11 |
| 1000 | 0.04 | 2.13 | 0.04 | 0.65 | 0.01 | 1.48 | 0.05 | 4.47 | 0.12 |

The fluid simulation makes predictions for the six Boolean logic gates (AND, NAND, OR, NOR, XOR, and XNOR). The output interpretation is based on a threshold calculated case per case on the average of the last 200 timesteps; Values over the threshold are considered “true/1”. During the first timesteps, the output produces not-distinguishable values. After ∼600 timesteps, each trend differentiates from the other, classifying the query pattern correctly. T-test is performed: p-values are significant among all the groups of values that are expected to be true or false depending on the context (p<0.0001) (Figure 5A). The mean and standard deviations of the last 200 timesteps’ output values for all the six logic gate tests are reported in Table 6.

**Table 6.**
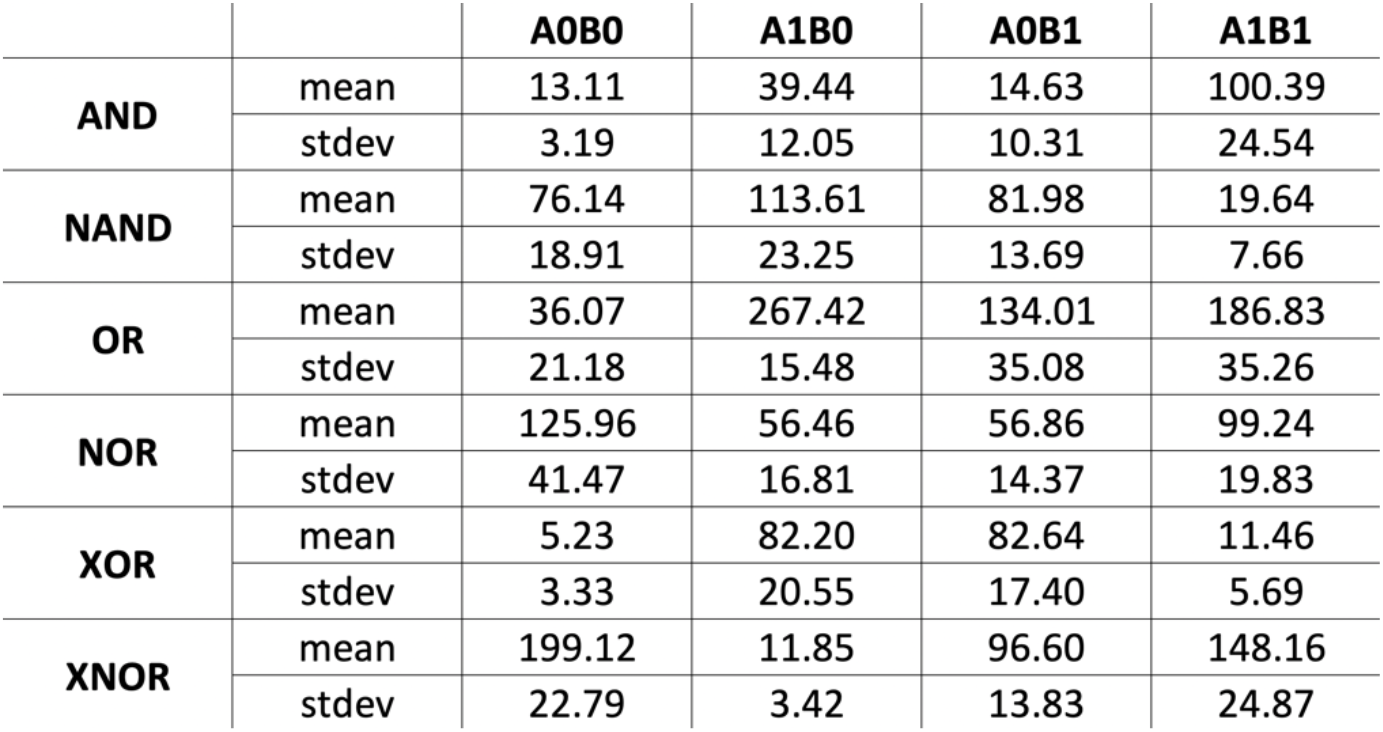
Mean and standard deviations of the output values registered in the last 200 timesteps for the six logic gate tests.

**Figure 5.**
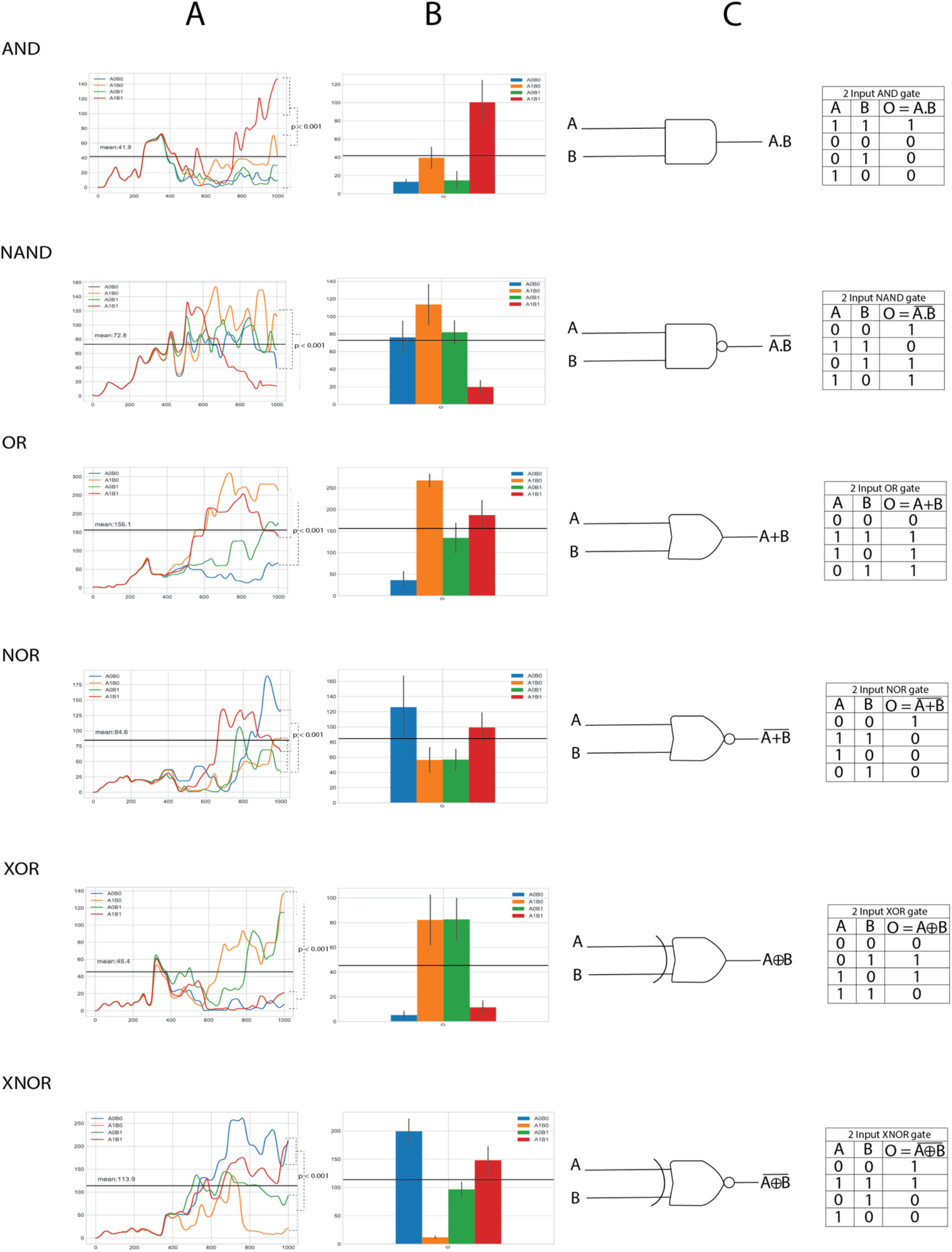
Tests on the six Boolean logic-gates (AND, NAND, OR, NOR, XOR, and XNOR). **A**: for each logic gate, the output’s velocity values are registered for all the test duration (1000 timesteps). Each color-coded line represents the trend corresponding to a tested pattern: blue A0B0, orange A1B0, green A0B1, and red A1B1. Initially, the output produces not-distinguishable values if compared to the others. The black threshold line is calculated based on the mean of the trends during the last 200 timesteps. After 600 timesteps, each trend differentiates, classifying the queried pattern correctly. The dashed lines report p-values by T-test (p<0.0001). **B:** Histograms represent the average of the last 200 timesteps’ output values, with relative standard deviations. **C**: schematical picture of logic gate and truth tables. The pattern’s order of the truth tables is the same used for training.

## 4 Discussion

The Equivalence explains how thermodynamics embodies the dissipative process that minimizes the internal entropy. This process leads to a continuous variation of all the particle’s coordinates. It goes ahead until each particle pair finds the cost minimum (the best minimum) for their potential function. I presume that the Gibbs free energy for this system matches the point in which the global error reaches its minimum. This condition may correspond to the moment in which the minimizing process produces ordered trajectories of particles, obtained in the solid and dense fluid phases (Figure 4D). The reduction of global potential energy also confirms the emergence of this order.

An interpretation of the phenomenon is that, in essence, for each particle’s pairs, the force’s vector 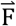 pushes the particles in the direction where the potential energy reduces. 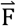 depends on spatial coordinates *(x, y)*, and the gradient −∇u(r) represents its logistic regression.

For the Equivalence, the gradient −∇u(r) of an LJ fluid corresponds to the cost function’s partial derivative (Table 1). For a Neural Network, the θ vector represents its weights (or in other terms, θ is the minimized configuration of all the neuron values after a training phase). Similarly, the corresponding θ vector is represented by the particle’s coordinates, velocities, and accelerations for the soft-disks fluid, dealing with minimizing the whole system’s structure after a completed training phase. Since that the fluid’s u(r), matches with the Neural Network’s J(θ), then:

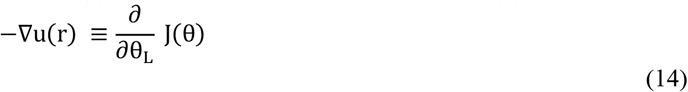

The soft-disks fluid behaves like a gradient descent for the LJ potential functions of all the particle couples of the fluid. The following integral equation expresses the relationship between a conservative force’s potential energy and the force itself. It is given by:

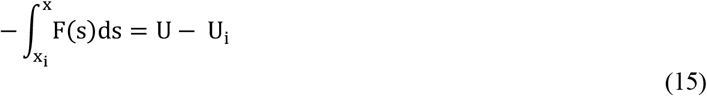

The Equivalence makes prerequisites for providing proofs of learning and evidence of discernment of a simulated fluid. The choice of a fluid model for this work is necessary for simplicity reasons. Indeed, the fluid approach is decisively more affordable than working on rigid molecules, in which more complex force fields govern interactions. Conversely, the fluid’s drawback is that the output is not kept after a certain number of timesteps. The output velocities tend to diverge, making it impossible to construct complex circuitries based on the combination of different logic gates. A possible explanation of this problem may be that fluid is a system in which particles are not tightly bound. Molecular Dynamics drives the evolution of the fluid by the variation of its potentials. The learning has to be considered part of an order that emerges as an adaptation in establishing equilibrium with energy and thermal conservation.

For the first time, this work demonstrates that a simulated chemical system can learn from examples. The learning is a consequence of a minimizing process among the particles and involves their velocities, accelerations, and forces. The minimizing process happens without the help of external actions. The gradient descent properties are embedded in the disposition of its particles.

One restriction is that the training is strongly limited to the patterns’ sequence presentation in the fluid’s molecular learning. If we change the order in which the patterns are presented, the output prediction could be affected. This limitation makes the soft-disks fluid an unconventional learning system. Instead, in classic machine learning, training success does happen regardless of the pattern’s chronological presentation. We assume this restriction is a fluid’s peculiarity dued to the strong dependence on the initial conditions.

This work aims to explain how a soft-disks fluid model uses its intrinsic minimizing properties in learning and predicting outputs. But it appears clear as a more careful interpretation of its results may lead to a series of considerations concerning a broader vision of Nature’s learning processes. The learning may belong to networks in which a gradient descent process exists, minimizing the real outputs’ error.

Belonging the learning’s properties to every kind of network, we face a natural process that could involve other types of molecules, like rigid biomolecules governed by different types of potentials, particularly receptors.

Thus, all the observations included in this work could have an impact on molecular medicine. For example, the effect of receptor desensitization may be seen as a receptor’s overtraining for its ligand.

From molecular complexes to biological organisms, the self-organization plays its role across all the elements; here, an emerging cognitive system, whose organization defines a domain of interactions, can act with relevance to the maintenance of itself (Maturana and Varela 1980). This interaction domain seems to be a sufficient condition, such as to be considered a cognitive system.

The work has highlighted the Lennard-Jones potential role in constructing a domain of interaction for a simple atoms system, like a fluid. The results refer to a simulated system far from being a real system. The simulation approach based on thermodynamics has revealed an interaction domain’s formation between all the particles.

The world of Molecular Computing and industry could benefit from the approaches here described, focusing on controllable chemical devices, programmable at the atomic level, and showing learning abilities at a molecular level. Maybe in the future, a deeper understanding of all the forces and potentials governing more complex molecules will lead to a significant comprehension of their intrinsic mechanism, suggesting new hints for a theory on real chemistry’s computational universality.

## Statistics

All the statistics are performed with GraphPad Prism 9.

## Software Availability

The algorithm comes as a Python 3.8 implementation. The code is available for download at https://github.com/lucazammataro/SDFL

## References

Adleman, L. M. (1994). “Molecular computation of solutions to combinatorial problems.” Science 266(5187): 1021–1024.

Goldt, S. and U. Seifert (2017). “Stochastic Thermodynamics of Learning.” Phys Rev Lett 118(1): 010601.

Gori, M., M. Maggini and A. Rossi (2016). “Neural network training as a dissipative process.” Neural Netw 81: 72–80.

Hylton, T. (2020). “Thermodynamic Neural Network.” Entropy 22(3).

Maturana, H. R. and F. J. Varela (1980). Autopoiesis and Cognition: The Realization of the Living, Springer Netherlands.

Nicolis, G. and I. Prigogine (1977). Self-organization in nonequilibrium systems : from dissipative structures to order through fluctuations, Wiley.

Rapaport, D. C. (2002). “Molecular dynamics simulation of polymer helix formation using rigid-link methods.” Phys Rev E Stat Nonlin Soft Matter Phys 66(1 Pt 1): 011906.

Rapaport, D. C. (2004). The art of molecular dynamics simulation. Cambridge, UK; New York, NY, Cambridge University Press.

Sienko, T. (2003). Molecular computing. Cambridge, Mass., MIT Press.

Virtanen, P., R. Gommers, T. E. Oliphant, M. Haberland, T. Reddy, D. Cournapeau, E. Burovski, P. Peterson, W. Weckesser, J. Bright, S. J. van der Walt, M. Brett, J. Wilson, K. J. Millman, N. Mayorov, A. R. J. Nelson, E. Jones, R. Kern, E. Larson, C. J. Carey, I. Polat, Y. Feng, E. W. Moore, J. VanderPlas, D. Laxalde, J. Perktold, R. Cimrman, I. Henriksen, E. A. Quintero, C. R. Harris, A. M. Archibald, A. H. Ribeiro, F. Pedregosa, P. van Mulbregt and C. SciPy (2020). “SciPy 1.0: fundamental algorithms for scientific computing in Python.” Nat Methods 17(3): 261–272.

Zammataro, L. (2020). “The Lennard-Jones potential: why the art of molecular dynamics is so fascinating, and why I got so emotionally overwhelmed.” https://towardsdatascience.com/the-lennard-jones-potential-35b2bae9446c.

